# DROSHA associates to DNA damage sites and is required for DNA repair

**DOI:** 10.1101/261289

**Authors:** Matteo Cabrini, Marco Roncador, Alessandro Galbiati, Lina Cipolla, Fabio Iannelli, Simone Sabbioneda, Fabrizio d’Adda di Fagagna, Sofia Francia

## Abstract

The DNA damage response (DDR) is the signaling cascade through which a cell recognizes DNA lesions, and promotes their resolution via the repair pathways of Non-Homologous End Joining (NHEJ), or Homologous Recombination (HR). We recently demonstrated that DROSHA boosts DDR signaling by processing damage-induced long non-coding RNAs into smaller DNA damage response RNAs (DDRNAs). However, the location at which DROSHA exerts its DDR functions, relative to sites of DNA damage, remains unknown.

To investigate DROSHA’s localization during DDR activation, we used the DiVA cellular system, which allows the controlled induction of several DNA double strand breaks (DSBs) in the human genome. Indeed, by genome wide chromatin immunoprecipitation followed by next generation sequencing, we demonstrate that DROSHA associates with DSBs. In support of this, DSB-recruitment of DROSHA is detectable at the single-cell level by Proximity Ligation Assay between DROSHA and known DDR markers, and by DNA damage in situ ligation followed by Proximity Ligation Assay (DI-PLA), which demonstrates proximity of DROSHA to DNA ends. DROSHA recruitment occurs at both genic and inter-genic DSBs, suggesting that its recruitment is independent from ongoing transcription preceding damage generation. DROSHA’s recruitment to DNA lesions occurs throughout the cell cycle, and with a preference for NHEJ-prone DSBs. Consistently, inhibition of the HR pathway increases DROSHA recruitment, and DROSHA knock down strongly impairs NHEJ efficiency in a GFP-reporter cellular system for monitoring NHEJ DNA repair. Overall, these results demonstrate that DROSHA acts locally at sites of DNA damage to promote NHEJ DNA repair.

## Introduction

DNA lesions continuously challenge genome integrity, and efficient DNA repair is crucial to avoid genome instability leading to cancer. The DNA damage response (DDR) is the cascade of events that detects, signals and repairs DNA lesions. DNA double-strand breaks (DSBs) represent one of the major activators of the pathway. These lesions are recognized by the MRE11/RAD50/NBS1 (MRN) sensor complex that recruits and activate the Ataxia telangectasia mutated protein kinase (ATM) at the site of the DNA lesion (Shiloh, 2006). This causes ATM autophosphorylation (pATM) as well as the local phosphorylation of the histone H2AX (γH2AX) that recruits additional ATM in a positive feedback loop. As a consequence, γH2AX spreads along the chromosome (Rogakou et al., 1999) and additional DDR factors, such as MDC1 and 53BP1, are recruited, leading to the formation of cytologically-detectable foci at damaged genomic loci (Ciccia and Elledge, 2010). From DDR foci the signal is transduced throughout the nucleus ultimately stimulating DNA repair, which can rely on either non-homologous end joining (NHEJ) or homologous recombination (HR) depending on the cell-cycle phase of the damaged cell and the chromosomal context in which the damage is induced. We have previously reported the existence of a novel class of DICER- and DROSHA-dependant small non-coding RNAs (sncRNAs), named DDRNAs (for DNA damage response RNAs), involved in local DDR activation (Francia et al., 2012), (Francia et al., 2016), (Francia, 2015), (d′Adda di Fagagna, 2014). DDRNAs bear the sequence of the damaged locus and enable the accumulation of 53BP1 and MDC1 (Francia et al., 2016) at chromatin surrounding the lesion thus promoting DDR foci formation (Francia et al., 2012).

More recently we demonstrated that DDRNA are processed from damage induced long non-coding RNA (dilncRNA) synthetized by RNA polymerase II (RNAPolII) starting from the exposed DNA ends of DSBs. Importantly, dilncRNA generation provides the RNA precursor for DDRNA generation upon DROSHA and DICER processing and, concomitantly, a sequence specific recruiting element for mature DDRNA together with DDR protein factors to which they associate (Michelini et al., 2017). Similarly, we have shown that DDRNAs with telomeric sequences activate DDR at unprotected telomeres, which resemble DSBs (Rossiello et al., 2017). Interestingly, DROSHA inactivation by siRNA prevents telomere fusions. Other groups have reported an involvement of sequence-specific DICER dependent ncRNA in DNA repair by HR (Gao et al., 2014), (Wei et al., 2012) and knockdown of DICER or DROSHA significantly reduced accumulation of two major HR factors, Rad51 and BRCA1 to DSBs (Wang and Goldstein, 2016).

So far, the emerging model suggests that DSBs stimulate local transcription by RNA pol II of long non-coding RNA molecules that following DROSHA and DICER processing are required for full DDR activation. Thus, both DDRNA biogenesis and function appear to occur locally. In agreement with this model, the presence of a phosphorylated form of DICER at DNA damage sites has been recently reported, where it mediates 53BP1 foci formation (Burger et al., 2017), (Burger and Gullerova, 2018).

Here we show that DROSHA associates with DSB occurring at endogenous sequences and its recruitment peaks within 5 kilobases from the DNA lesion. DROSHA association with DSB is robustly observed genome wide and at individual damaged sites and at single-cell level by Proximity Ligation Assay and (PLA) and DNA damage in situ ligation followed by PLA (DI-PLA), which specifically detect protein-protein and protein-DNA ends proximity, respectively. DROSHA accumulation at DNA ends does not seem to depend on pre-existing transcription and it occurs in all phases of cell cycle, both at DSBs prone to engage NHEJ and HR mediated-repair mechanisms. Functionally, DROSHA recruitment to damage sites appears to be relevant for DNA repair efficiency by NHEJ.

## Results

### DROSHA is recruited to DSBs

In the light of the involvement of DROSHA in DDR signaling, its predominantly nuclear localization and of its reported co-transcriptional role in the context of microRNA processing (Morlando et al., 2012), (Gromak et al., 2013), we wondered if this endoribonuclase is recruited to sites of DNA damage to locally process newly synthetized dilncRNA into DDRNAs. To test this hypothesis, we took advantage of the DIvA cellular system (for <u>D</u>SB inducible <u>v</u>ia <u>A</u>siSI), (Iacovoni et al., 2010), (Aymard et al., 2014), (Iannelli et al., 2017), (Aymard et al., 2017) a clonal U2OS cell line that stably expresses the AsiSI restriction enzyme fused with a modified oestrogen receptor ligand binding domain. Treatment of cells with 4-OH-Tamoxifen (4-OHT) triggers nuclear localization of the AsiSI enzyme and the generation of hundreds of sequence-specific DSBs uniformly distributed in the genome. This system offers many advantages compared to other DNA damaging treatments because it allows the generation of a homogeneous type of DNA lesions at sequence-specific sites in different contexts of the human genome (Iannelli et al., 2017). In addition, since each AsiSI consensus sequence can be unambiguously mapped on the human genome, this system permits the use of Chromatin immunoprecipitation (ChIP) followed by either next generation sequencing (ChIP-seq) or qPCR (ChIP-qPCR) analysis as an effective and unbiased method for identifying protein association with sites of DNA damage.

Since the AsiSI enzyme is sensitive to CpG methylation and to the structure of surrounding chromatin (Iacovoni et al., 2010) only a subset of all the potential AsiSi restriction sites present in the human genome are efficiently cut (Iannelli et al., 2017).

To precisely determine which AsiSI sites are efficiently targeted by the enzyme in the cell population studied, we performed ChIP-seq for γH2AX, a reliable marker of DNA damage. We identified a list of 361 sites that are cut by AsiSI as indicated by the significant enrichment in γH2AX signal in damaged cells respect to the undamaged condition and we decided to focus our analyses on the top fifty (top 50) sites, which are the subset of sites more robustly cut in the cell population and among independent experiments.

As previously reported in literature (Aymard et al., 2014), γH2AX ChIP-seq signal showed that, upon 4OHT induction, phosphorylation of histone H2AX established a megabase-large zone of modified chromatin surrounding the AsiSI-DSB (Figure 1A and B). Interestingly, parallel DROSHA ChIP-seq analyses indicated that this protein is robustly recruited to DNA damage loci and its accumulation sharply peaks around the AsiSI-induced break point (Figure 1C and D). In accordance with the average DROSHA profile, a closer inspection of the ChIP-seq pattern at individual DSBs (DSB I and DSB II detected as cut in (Iannelli et al., 2017)) confirmed the strong enrichment of DROSHA at DNA DSBs (Figure 1E). To better address whether the DSB genomic context could affect DROSHA recruitment, we defined and characterized three subgroups of cut AsiSI sites according to their localization at i)promoters, ii) coding sequences (CDS) or iii) intergenic regions (away from any annotated genetic unit) (Figure 1F). Importantly, DROSHA was detectable at all the AsiSI induced DSBs analyzed indicating that pre-existing transcription is not a prerequisite for DROSHA association with damaged chromatin (Figure 1F). Nevertheless, it is worth noticing that intergenic sites showed clear but somehow lower accumulation of DROSHA, despite comparable cut efficiency as measured by γH2AX accumulation, suggesting that transcription-associated chromatin state might stabilize DROSHA recruitment (Figure 1F).

**Figure 1.**
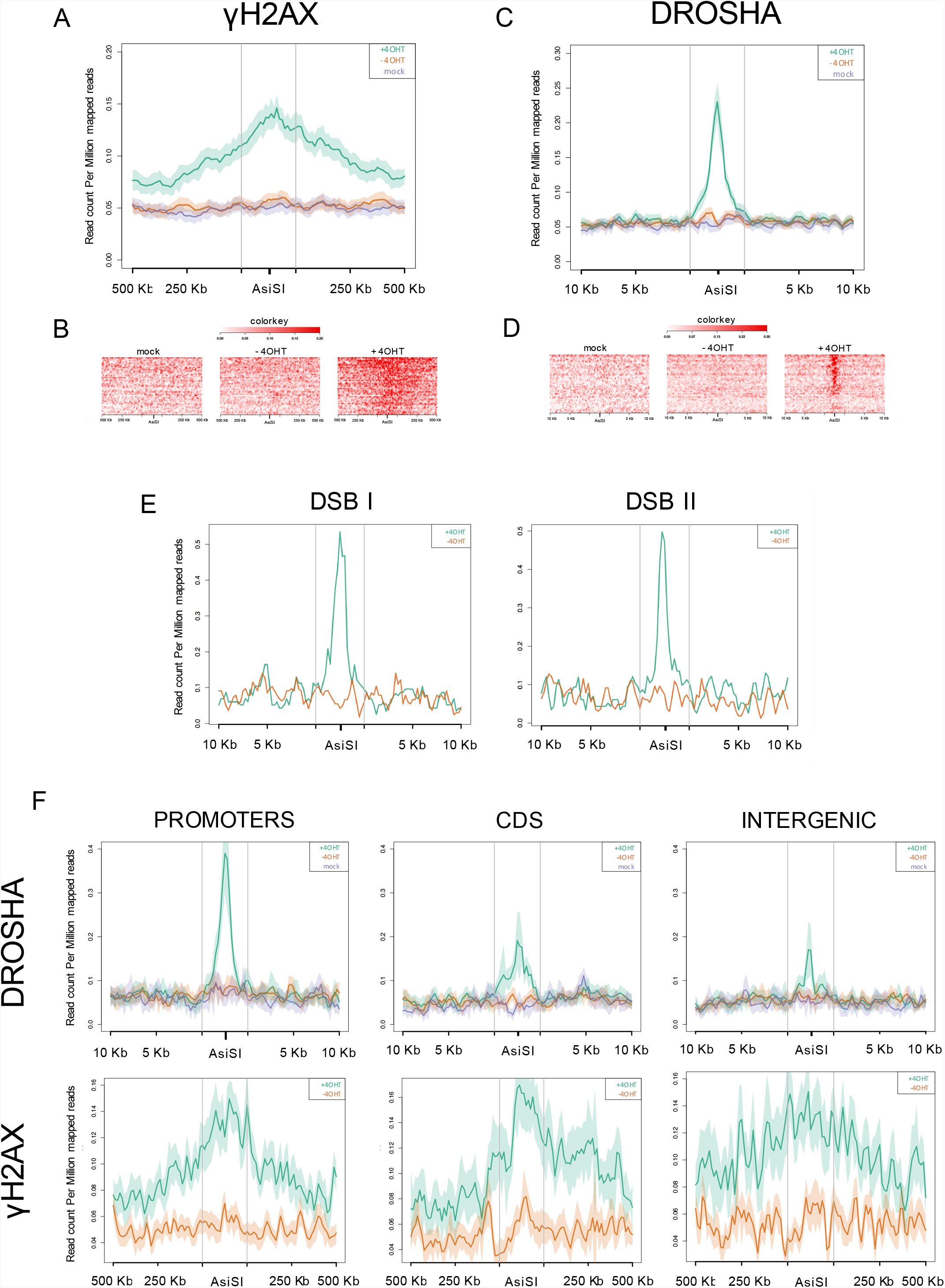
DROSHA accumulates to sites of DNA damage as determined by genome wide ChIP-seq. **A)**Coverage profile plot representing the RPM of averaged γH2AX ChIP-seq signal of the top 50 AsiSI sites, according to γH2AX ranking, over 1 Mb windows and centered at the AsiSI site, are shown for uninduced (red), induced (green) DIvA cells or mock (magenta) sample. **B)** Heatmap representation of γH2AX ChIP-seq signal across the 50 AsiSI sites shown in the plot above, sorted by γH2AX decreasing signal, over a 1 Mb window. **C)** Coverage profile plot representing the RPM of averaged DROSHA ChIP-seq signal of the top50 AsiSI sites, according to γH2AX ranking, over 20 Kb windows and centered at the AsiSI site, are shown for uninduced (red), induced (green) DIvA cells or mock (magenta) sample. **D)** Heatmap representation of DROSHA ChIP-seq signal across the 50 AsiSI sites shown in the plot above, sorted by DROSHA decreasing signal over a 20 Kb window. **E)** Coverage profile plot representing the RPM of DROSHA ChIP-seq signal in uninduced (red) and induced (green) samples of two representative AsiSI sites ranked in the top positions, according to γH2AX ranking. **F)** Coverage profile plot representing the RPM of averaged DROSHA and γH2AX signal of 8 AsiSI sites positioned in a region annotated as a Promoter, CDS or Intergenic, over 20 Kb windows and centered at the AsiSI site, are shown for uninduced (red), induced (green) DIvA cells or mock (magenta) sample. All the AsiSI sites studied are included in the top50 according to the γH2AX ranking.

To independently validate these genome wide conclusions in a distinct biological replicate at an individual site we performed ChIP-qPCR analysis at a representative AsiSI induced DSB. This analysis confirmed that both γH2AX and DROSHA levels increased significantly in the induced samples compared to the uninduced one (Figure 2A). Importantly, DROSHA knock down dramatically reduced the enrichment of DROSHA at DNA damage site, without altering γH2AX accumulation in the same sites, an observation consistent with our previous reports (Francia et al., 2012), (Francia et al., 2016), (Michelini et al., 2017) (Figure 2A and B). Importantly, these results also demonstrate the specificity of the antibody used. As a control, neither γH2AX nor DROSHA accumulate at a distal uncut region (Figure 2A). In addition, we used ChIP-qPCR to investigate the spreading of DROSHA from DSBs, using primers mapping at increasing distances (60bp, 1Kb and 2,5Kb) from the DNA damage site. This analysis revealed that, differently from γH2AX, DROSHA enrichment peaks in the vicinity of the DSB (within 2,5 Kb), suggesting that DROSHA might not depend on γH2AX histone modification for its localization at site of DNA damage (Figure 2C). Of note, DROSHA silencing causes a loss of signal for DROSHA in the vicinity of the break and also an increase of γH2AX accumulation consistent with a potentially reduced repair (Figure 2C).

**Figure 2.**
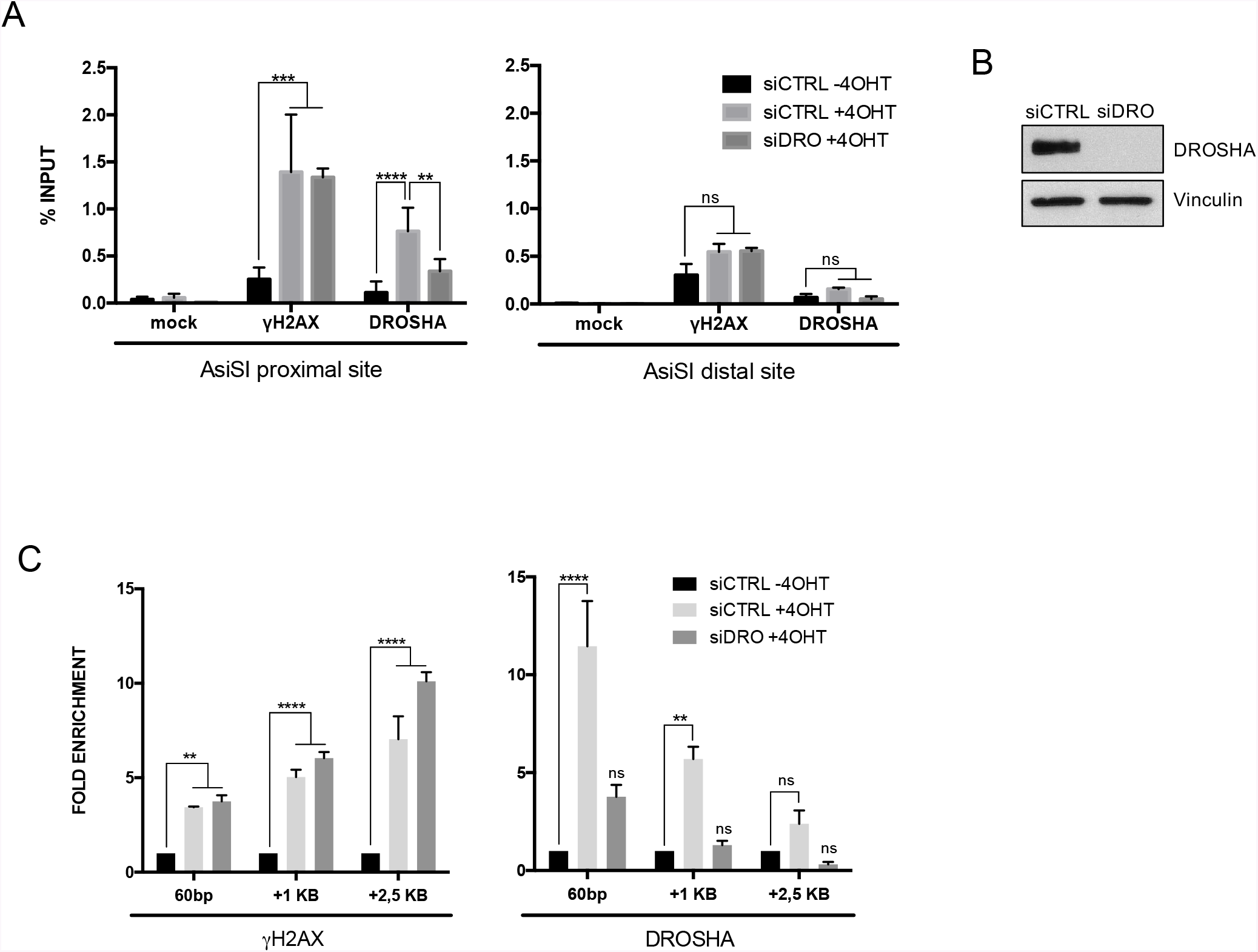
DROSHA is recruited close to the sites of DNA damage as detected by qPCR at individual AsiSI sites. **A)**The bar plot shows the percentage of enrichment, relative to the input, of mock (no antibody), γH2AX and DROSHA associated with genomic DNA, as detected by ChIP-qPCR in uninduced (-4OHT) or induced (+4OHT) DIvA cells with or without DROSHA knock down, with primers matching DSB I (proximal site) or a genomic region far from any annotated AsiSI sites (distal site). Error bars indicate standard deviation among 4 independent experiments. P-values were calculated by one-way ANOVA with multiple comparison. **B)**DIvA cells were transfected with control siRNA (siCTRL) or siRNA against DROSHA (siDRO). 72h later, knock down efficiency was evaluated by western blotting. Vinculin was used as loading control. **C)** The bar plot shows the percentage of enrichment, relative to the input, of mock (no antibody), γH2AX and DROSHA associated with genomic DNA, as detected by ChIP-qPCR in uninduced (-4OHT) or induced (+4OHT) DIvA cells with or without DROSHA knock down, with primers matching DSB II (AsiSI-DSB), 1Kb or 2,5Kb far from DSB II (representative experiment, n=2). P-values were calculated by one-way ANOVA with multiple comparison.

Next, we attempted to detect DROSHA recruitment to DNA damage sites by standard immunofluorescence. Indeed, recruitment of DDR proteins to DNA lesions can be directly visualized by immunofluorescence in the form of cytological detectable foci. Immunostaining for DROSHA in a cellular reporter system that employs a mCherry-LacI-FokI nuclease fusion protein to create multiple DSBs within a single genomic locus containing Lac operone repeats (Shanbhag et al., 2010) failed to show a detectable accumulation of the protein at the damaged locus despite a clear focal accumulation of γH2AX (Figure 3A). However, it is known that some DDR factors, such as KU70/80 and DNAPKcs, which are recruited to DNA ends with high specificity but do not spread on the chromatin flanking the DSB, cannot be efficiently detected as DNA damage foci by immunofluorescence (Britton et al., 2013; Polo and Jackson, 2011) (Britton et al., 2013). Thus, we sought to assess DROSHA accumulation at individual sites of DNA damage in single cells by performing PLA with antibodies against DROSHA and γH2AX. This approach allows to detect proximity among proteins within 40 nm. PLA revealed several nuclear fluorescent signals following DNA damage generation. These signals were specific as demonstrated by their loss upon DROSHA knockdown, which does not reduce H2AX phosphorylation (Figure 3B, C and D). To strengthen these observations, we took advantage of a novel method recently developed in our laboratory, named ‘DNA damage in situ ligation followed by Proximity Ligation Assay’ (DI-PLA) (Galbiati et al., 2017). This technology allows the study of the recruitment of a given protein at DNA ends. Briefly, in fixed and permeabilized cells, DNA ends are blunted and ligated to a biotinylated oligonucleotide (Crosetto et al., 2013) which permanently tags DSB-ends; then PLA is performed using primary antibodies against biotin and a partner antibody against the protein of interest, DROSHA in this case (see cartoon in Figure 3E). By this approach, we observed a significant increase of DI-PLA signals upon DNA damage generation, indicating DROSHA presence in close proximity to DNA ends of DSBs (Figure 3F and G). Again, the signal observed was specific since DROSHA knockdown dramatically reduce it (Figure 3F, G and H). Therefore, our ChIP data together with these single-cells observations indicate that upon DSBs generation DROSHA is recruited to DNA damage sites and its accumulation is concentrated around the break point.

**Figure 3.**
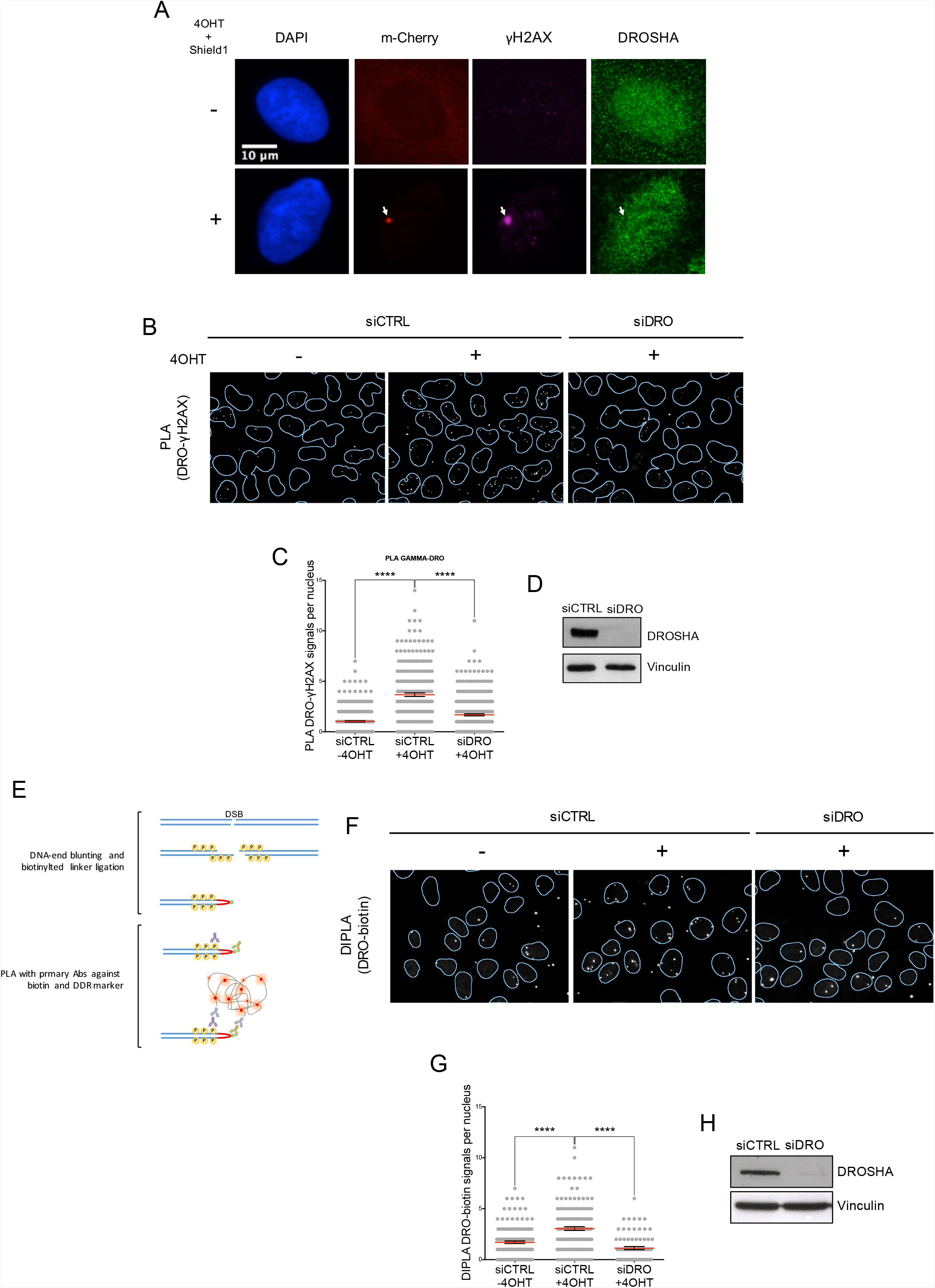
PLA and DIPLA reveal γH2AX-DROSHA proximity. **A)**U2OS-DSB reporter cells expressing mCherry-LacI-FokI nuclease fusion protein, were co-stained with DAPI, γH2AX and DROSHA, before and after 4OHT and Shield1 treatment to induce cut (4 h). White arrow indicates FokI-cut locus. **B)**Representative images of PLA between γH2AX and DROSHA in uninduced (-4OHT) and induced (+4OHT) DIvA cells with or without DROSHA knock down. **C)** The scatter plot represents the number of nuclear PLA signals as measured by Cell Profiler software. The red bar represents the mean ± SEM (300 cells, n=2). P-values were calculated by one-way ANOVA with multiple comparison. **D)** DIvA cells were transfected with control siRNA (siCTRL) or siRNA against DROSHA (siDRO). 72h later, knock down efficiency was evaluated by western blotting. Vinculin was used as loading control. **E)** Schematic cartoon of DI-PLA. **F)** Representative images of DI-PLA between biotin and DROSHA in uninduced (-4OHT) and induced (+4OHT) DIvA cells with or without DROSHA knock down. **G)**The scatter plot represents the number of nuclear DIPLA signals as measured by Cell Profiler software. The red bar represents the mean ± SEM (150 cells, n=2). P-values were calculated by one-way ANOVA with multiple comparison. **H)**DIvA cells were transfected with control siRNA (siCTRL) or siRNA against DROSHA (siDRO). 72h later, knock down efficiency was evaluated by western blotting. Vinculin was used as loading control.

### DROSHA is recruited at DSBs throughout the cell-cycle

We have previously reported a role for DROSHA in DDR signaling (Francia et al., 2012), (Francia et al., 2016), (Michelini et al., 2017), (Rossiello et al., 2017), (Pignataro et al., 2017), however its contribution to NHEJ, the main DSB repair pathway in mammalian cells, has not been yet clarified.

NHEJ is active during all the stages of the cell cycle while HR is restricted to S and G2 phases, when homolog sister chromatids are present (Hustedt and Durocher, 2016). Thus, we tested whether DROSHA recruitment to sites of DNA damage occurs throughout the cell cycle or preferentially in S-G2. To this aim we took advantage of the FUCCI cell system (Goto et al., 2015) a fluorescent protein-based system that employs both a red (RFP) and a green (GFP) fluorescent protein fused to different regulators of the cell cycle to allow a direct visual readout of cell cycle phase for every cell in the population (Figure 4A). Upon treatment with the radiomimetic drug neocarzinostatin (NCS), we assessed DROSHA localization to DNA ends by PLA between γH2AX and DROSHA (Figure 4B) in FUCCI cells. Automated quantification of nuclear PLA signals revealed no differences in DROSHA-γH2AX proximity among the G1 or S-G2 sub-populations of cells (Figure 4 C). This observation suggests that the recruitment of DROSHA to DSBs is not restricted to cells in S/G2, as expected for an HR-specific factor, but it is an event that could be required for both NHEJ and HR.

**Figure 4.**
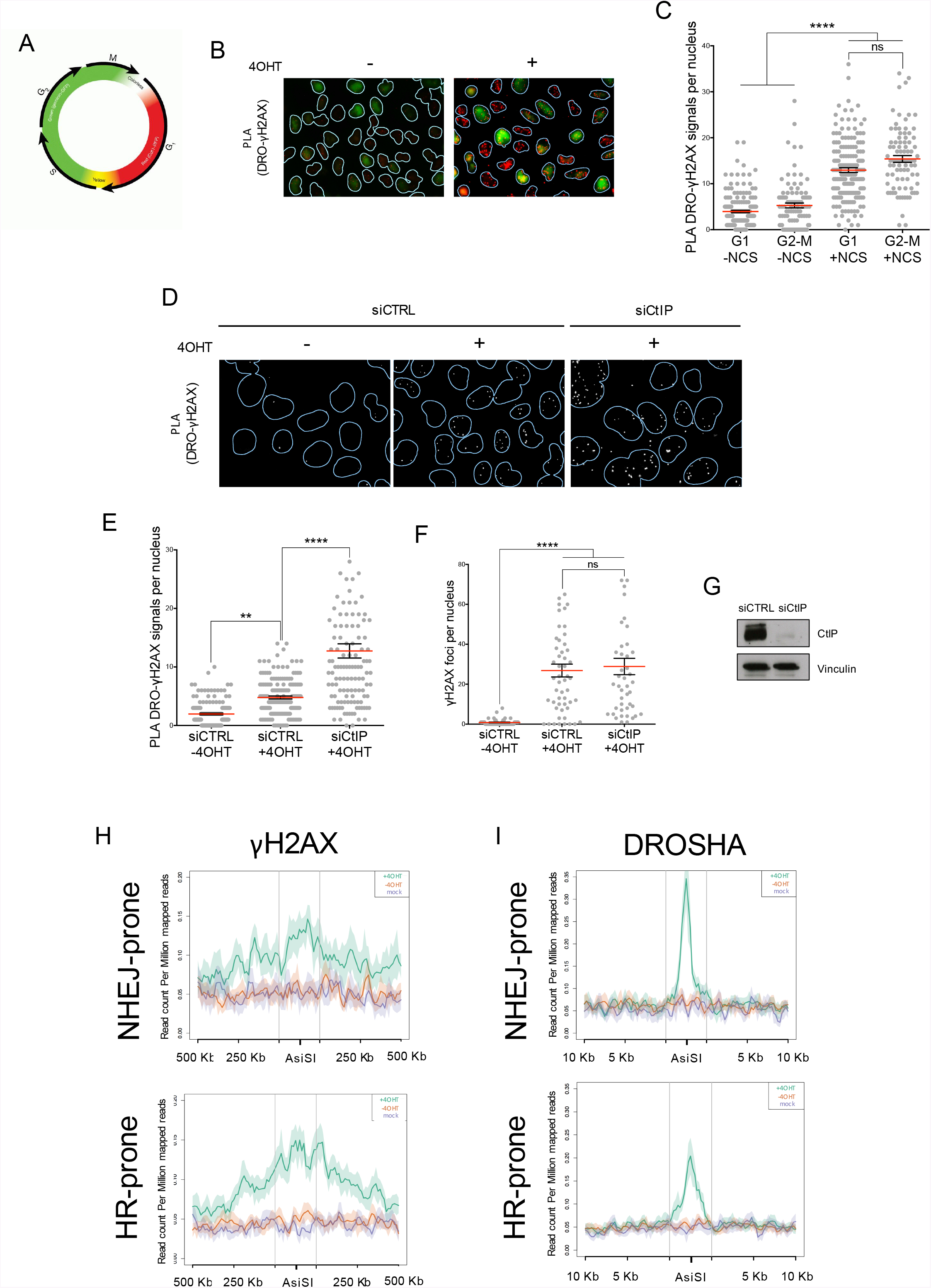
DROSHA recruitment to DSBs occurs throughout the cell cycle and preferentially at NHEJ-prone DSBs. **A)** Schematic cartoon of the the FUCCI (Fluorescence Ubiquitination Cell Cycle Indicator) system which labels cells in G1 phase red and cells in G2-M phases in green. (picture from https://www.thermofisher.com). **B)** Representative images of PLA between γH2AX and DROSHA in untreated and NCS-treated (12ng/ml) HeLa-FUCCI cells. **C)** The scatter plot represents the number of nuclear PLA signals as measured by Cell Profiler software. The red bar represents the mean ± SEM (250 cells, n=2). P-values were calculated by one-way ANOVA with multiple comparison. **D)** Representative images of PLA between γH2AX and DROSHA in uninduced (-4OHT) and induced (+4OHT) DIvA cells with or without CtIP knock down. **E)** The scatter plot represents the number of nuclear PLA signals as measured by Cell Profiler software. The red bar represents the mean ± SEM (150 cells, n=2). P-values were calculated by one-way ANOVA with multiple comparison. **F)** The scatter plot represents the number of γH2AX foci evaluated by immunofluorescence, and measured by Cell Profiler software. The red bar represents the mean ± SEM (100 cells, n=1). P-values were calculated by one-way ANOVA with multiple comparison. **G)** DIvA cells were transfected with control siRNA (siCTRL) or siRNA against CtIP (siCtIP). 72h post transfection, knock down efficiency was evaluated by western blotting. Vinculin was used as loading control. **H)** Coverage profile plots representing the RPM of averaged γH2AX ChIP-seq signal of the NHEJ-prone (upper panel) and HR-prone (lower panel) sites, over 1 Mb windows and centered at the AsiSI site, are shown for uninduced (red), induced (green) DIvA cells or mock (magenta) sample. **I)** Coverage profile plots representing the RPM of averaged DROSHA ChIP-seq signal of the NHEJ-prone (upper panel) and HR-prone (lower panel) sites, over 20 Kb windows and centered at the AsiSI site, are shown for uninduced (red), induced (green) DIvA cells or

### DROSHA recruitment to DSBs is enhanced by HR suppression and occurs preferentially at NHEJ-prone DSBs

To investigate the DNA repair pathway involving DROSHA, we tested the impact of DROSHA recruitment to DSBs upon HR suppression. To this aim, we prevented HR initiation by knocking down CtIP, an essential HR factor involved in DNA end resection, forcing cells toward NHEJ repair choice (Yun and Hiom, 2009). Excitingly, we observed that HR inhibition enhanced the number of γH2AX-DROSHA PLA signals per nucleus (Figure 4D, E and G). Importantly, the number of γH2AX foci did not increase upon CtIP silencing (Figure 4F) ruling-out the possibility that the augmented PLA signal observed depended on a global increase of DNA damage.

A previous study (Aymard et al., 2014) in the DIvA cellular system reported that in G1 the recruitment of the NHEJ factor XRCC4 occurs at all AsiSI induced DSBs, while in G2 RAD51 is recruited at a subset of DSBs. On the basis of RAD51/XRCC4 ratios, this analysis highlighted two subgroups of AsiSI induced DSBs referred as HR-prone and NHEJ-prone sites (Aymard et al., 2014). Thus, we decided to exploit this information to further validate a link between DROSHA and the NHEJ repair pathway. We subdivided our 50 most cut AsiSI induced DSBs, in HR- or NHEJ-prone. Next, we plotted the average profile of γH2AX and DROSHA for each subgroup. Despite exhibiting a robust and comparable γH2AX induction, the two subset displayed differences in DROSHA abundance (Figure 4H and I). Consistently with previous results, DROSHA enrichment was higher in the NHEJ-prone subgroup compared to the HR prone one (Figure 4I). Taken together these observations reveal a stronger affinity of DROSHA for DSB which are engaged in NHEJ repair pathway.

### DROSHA controls DNA repair by NHEJ

To assess the functional relevance of the observed preferential recruitment of DROSHA to NHEJ-prone AsiSI sites, we exploited the U2OS EJ5 cellular system, a well-established GFP-based reporter cell system in which the reconstitution of a functional GFP gene occurs after DNA damage generation and repair by NHEJ repair (Gunn et al., 2011), (Gunn and Stark, 2012) (see cartoon in Figure 5A). Interestingly, we observed that the number of GFP-positive cells was significantly reduced in U2OS EJ5 cells knocked down for DROSHA, to an extent similar to that observed upon KU80 knockdown, a central player in NHEJ (Figure 5B, C and D). Importantly, in this cell line, cell-cycle distribution is not altered by DROSHA inactivation (Figure 5E).

**Figure 5.**
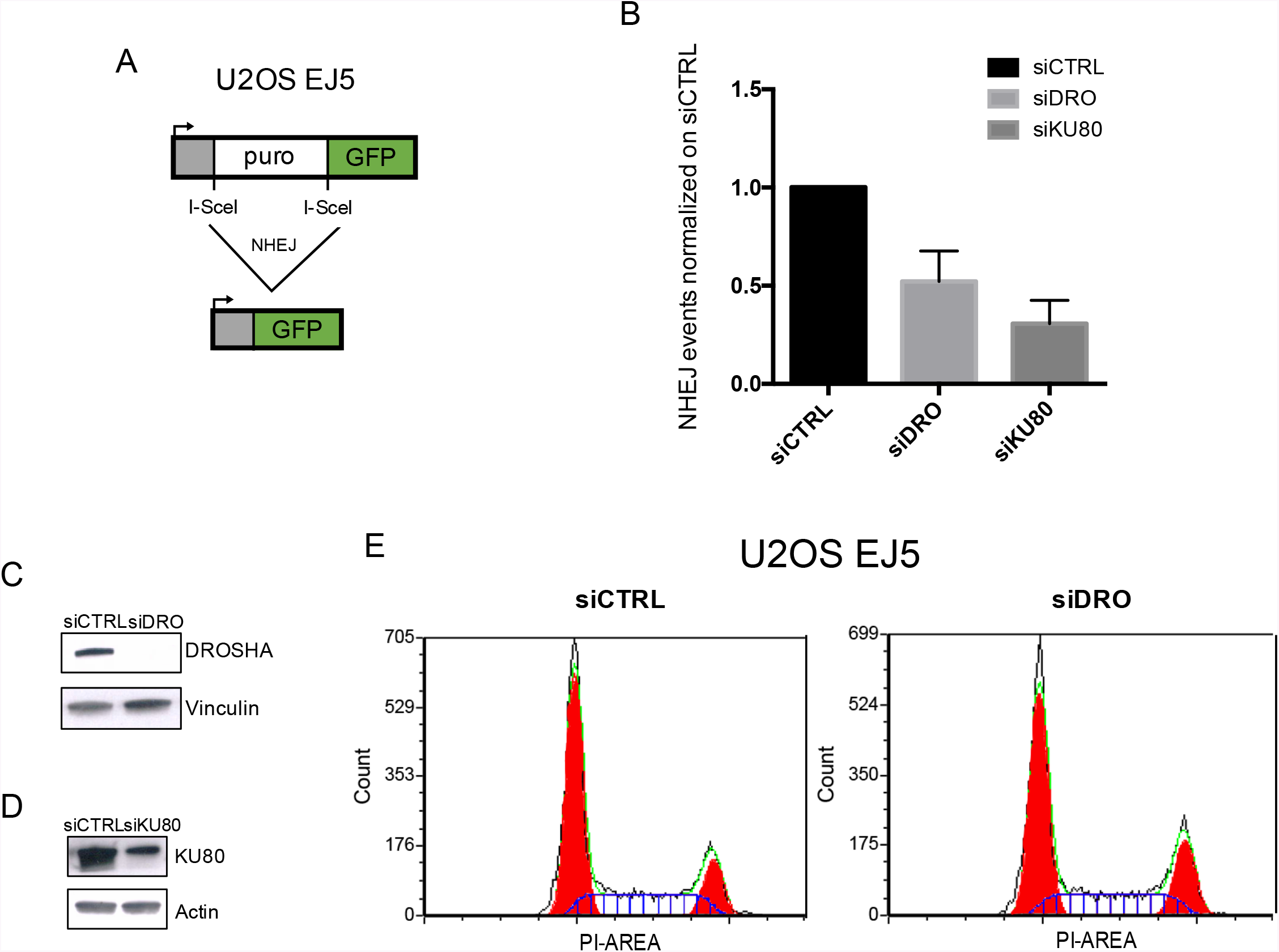
DROSHA knock down impairs DNA repair by non-homologous end joining. **A)** Schematic cartoon of the EJ5-GFP reporter used to monitor non-homologous end joining (NHEJ) in U2OS cells. **B)** The bar plot represents the fold enrichment of GFP-positive cells, normalized on the siCTRL ISceI-cut sample, in U2OS EJ5 cells transfected with control siRNA (siCTRL), siRNA against DROSHA (siDRO) or siRNA against KU80 (siKU80) and transfected with ISceI plasmid to induce the cut. Error bars indicate standard deviation among 3 independent experiments. **C and D)** U2OS EJ5 cells were transfected with control siRNA (siCTRL) and siRNA against DROSHA (siDRO) **(C)** or siRNA against KU80 (siKU80) **(D)**. 72h later, KD efficiency was evaluated by western blotting. Vinculin or Actin were used as loading control. **E)** FACS profiles of U2OS EJ5 cells transfected with control siRNA (siCTRL) and siRNA against DROSHA (siDRO).

Taken together, these results demonstrate an unanticipated role for DROSHA at sites of DNA damage in promoting repair by NHEJ.

## Discussion

Previously our group and others demonstrated that DNA damage sites are not transcriptionally silent (Michelini et al., 2017); (Capozzo et al., 2017). On the contrary, they induce the synthesis of non coding transcripts, which can be further processed into smaller RNA by components of the RNA interference machinery and together play a supportive role in DDR signaling (Francia et al., 2012), (Wei et al., 2012), (Michalik et al., 2012), (Michelini et al., 2017). We recently reported that RNAPII is recruited to DSBs in a MRN-dependent manner, where it synthesizes dilncRNAs from and towards DNA ends. Induction of dilncRNA was observed in several distinct cell systems, including U2OS DIvA cells, and accumulation of these transcripts has been reported following DROSHA knockdown (Michelini et al., 2017). Thus, DROSHA may act immediately upon dilncRNA synthesis, to generate an intermediate non-coding RNA product amenable for DICER enzymatic processing. However, whether non-coding RNA processing occurs locally at sites of DNA damage in the nucleoplasm or in the cytoplasm of damaged cells, remained to be addressed. A debate about the possibility that RNA interference factors might be active in the nucleus, was also exacerbated by the fact that DICER has been first shown to be exclusively cytoplasmatic (Much et al., 2016) and then to be recruited to sites of DNA damage in its phosphorylated form (Burger et al., 2017), (Burger and Gullerova, 2018).

Here we report for the first time that DROSHA accumulates at sites of DNA damage. DROSHA association with DNA lesions was assessed both by genome wide ChIP followed by next generation sequencing and by ChIP-qPCR at individual DSB, as well as by imaging techniques that allow the visualization of its recruitment to individual DSB at the single-cell level.

DROSHA recruitment appears as an event shared among DSBs generated in different genomic context (including promoters, CDS and intergenic regions) and it occurs throughout the cell cycle. In addition, its recruitment is restricted to the first kilobases near the DNA end, a localization which very much resembles that of NHEJ-repair factors such as XRCC4 (Aymard et al., 2014). Consistently, DROSHA accumulates preferentially at NHEJ-prone sites and its association with damaged chromatin is enhanced by NHEJ stimulation and HR suppression, as achieved by CtIP silencing. Indeed, DROSHA knockdown strongly impairs NHEJ repair efficiency to an extent similar to the inactivation of KU80, a central player in this repair pathway. Consistent with this result, a recent report from our group demonstrated that DROSHA inactivation results in a decrease in telomeric fusions, a process that relies on NHEJ (Rossiello et al., 2017).

Concomitantly, some reports suggested a link between DNA damage-dependent small RNA and the HR repair pathway and it has been recently observed that knockdown of DICER or DROSHA attenuates Rad51 and BRCA1 foci formation, two key players in HR (Wang and Goldstein, 2016).

Overall, it is becoming more and more apparent that DICER and DROSHA are essential for DDR activation and are involved in DNA repair (d′Adda di Fagagna, 2014), (Francia, 2015), a function that might prevent cancer development. Indeed, emerging evidence shows that altered expression or mutations of both DICER and DROSHA predispose to cancer (Lin and Gregory, 2015). Somatic mutations of DROSHA have been shown to be frequent and underlie high risk of Wilms tumors (Torrezan et al., 2014), (Wegert et al., 2015), (Spreafico et al., 2016) and DROSHA depletion has been reported to be implicated in the promotion of a migratory phenotype in lung cancer cells (Frixa et al., 2017). Moreover, expression of DROSHA has been recently shown to be impaired in breast cancer patients, although the molecular mechanisms by which this may be relevant for cancer development remain to be defined (Poursadegh Zonouzi et al., 2017).

In conclusion, we have identified an unanticipated role of DROSHA at sites of DNA damage which is not restricted to DDR signaling but it is relevant also for DNA repair. It is important to mention that NHEJ repair events are the most frequently occurring in our body, especially in terminally differentiated cells such as neurons, with important implications in the understanding of molecular mechanism behind neurodegeneration. DROSHA has been recently involved in the cellular response to DNA damage in neuronal cells carrying a Parkinson disease-linked mitochondrial mutation (Pignataro et al., 2017).

## Matherial and Methods

### Cell culture and treatments

DIvA cells (AsiSI-ER-U20S) (Iacovoni et al., 2010) were cultured in Dulbecco’s modified Eagle’s medium (DMEM) (Life-Technologies) w/o phenol red supplemented with 10% fetal bovine serum (FBS) (Euroclone), 1% L-Glutamine, 1% pyruvate, 2.5% HEPES and 1% penicillin/streptomycin. Cells were selected with puromycin (1 μg/ml). For AsiSI-dependent DSBs induction, cells were treated with 300 nM 4OHT (Sigma-Aldrich) for 4 hours. The U2OS-DSB reporter cell line developed by Greenberg Laboratory (Shanbhag et al., 2010) was grown in DMEM (Lonza) supplemented with 10% FBS Tetracycline tested (Euroclone), 1% L-glutamine, 1% penicillin/streptomycin. Cells were selected with puromicin (2 μg /mL) and G418 (400 μg/ml). DSBs induction was achived by the addition of 1 μM Shield-1 (Clontech) and 1 μM 4OHT for 4 h.HeLa-FUCCI cells were grown in DMEM (Euroclone), supplemented with 10% FBS (Euroclone), 1% L-Glutamine and 1% penicillin/streptomycin. DNA damage was induced using the radiomimetic drug Neocarzinostatin (NCS) (Sigma- Aldrich) at a concentration of 12 ng/ml for 20 min at 37°C.

All the cells lines were grown at 37°C under a humidified atmosphere with 5% CO2.

### Antibodies

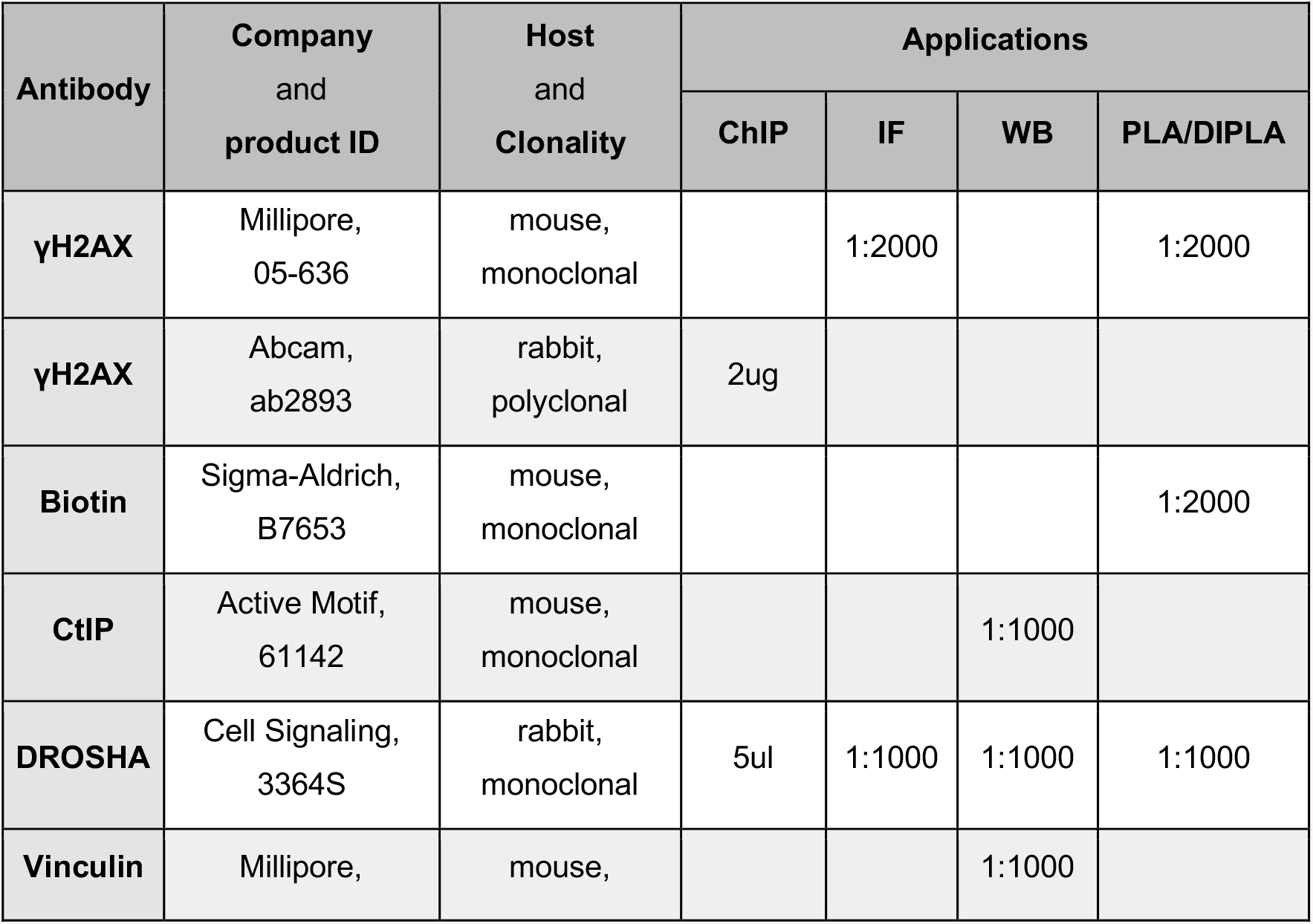
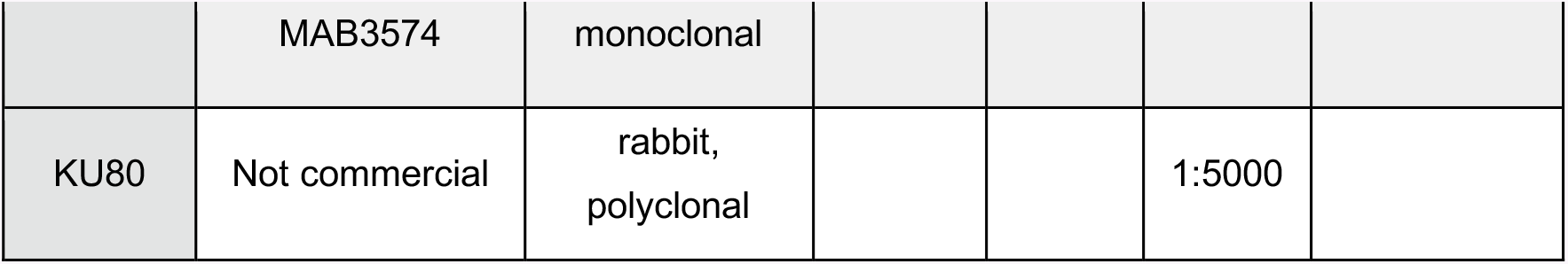

### Chromatin Immunoprecipitation

Cells were cross-linked for 5.5 min at room temperature with Fixation Buffer (1% formaldehyde; 100 mM NaCl; 1 mM EDTA; 0.5 mM EGTA; 50 mM HEPES pH 7.4). Cross-linking was quenched by addition of glycine (125 mM). Fixed cells were rinsed twice in 1X PBS, collected by scraping and centrifuged at 2000 rpm for 5 min at 4°C. Pellets were re-suspended in cold B1 Buffer (0.25% Triton X-100; 1 mM EDTA; 0.5 mM EGTA; 10 mM Tris pH 8; Proteases inhibitors (Roche); Microcystin (Enzo Life Sciences)) by mixing for 10 min on a rotating wheel at 4°C and then centrifuged at 2000 rpm for 5 min at 4°C. The same steps were repeated with cold Buffer B2 (200 mM NaCl; 1 mM EDTA; 0.5 mM EGTA; 10 mM Tris pH 8; Proteases inhibitors (Roche); Microcystin (Enzo Life Sciences)). Finally, pellets were re-suspended in cold Buffer B3 (TE 1X; EGTA 0.5 mM) in a suitable volume. Pellets were sonicated using a Focused-Ultrasonicator Covaris (duty: 5.0; PIP: 140; cycles: 200; amplitude: 0; velocity: 0; dwell: 0; microTUBEs with AFA fiber). Sonicated chromatin was diluted in RIPA buffer (1% TritonX-100; 0,1% Na-Deoxycholate; 0,1% SDS; 64 mM NaCl; 10 mM Tris HCl pH 8.0) to give a concentration of approximately 100 μg in 400 μl per ChIP. Samples were pre-cleared for 2 h, rotating at 4°C, with 20 μl of magnetic beads (Dynabeads^®^ Protein G, LifeTechnologies) per ChIP. Sample were then incubated overnight rotating at 4°C with specific antibodies (see Antibodies section for a complete list) or no antibody (mock). The bound material was recovered by 2 h incubation with 20 μl of magnetic beads per ChIP. Beads were then washed, rotating at 4°C for 10 min, four times in RIPA buffer, once in LiCl buffer (250 mM LiCl; 0.5% NP-40; 0.5% Na Deoxycholate; 1 mM EDTA; 10 mM Tris-HCl pH 8) and finally in 1X TE. ChIPed material was eluted by 15 min incubation at 65°C with 150 μl Elution Buffer (1% SDS; 10 mM EDTA; 50 mM Tris HCl pH 8). Samples were reverse-crosslinked by incubation with proteinase K (Invitrogen) at 37°C for 5 h and then at 65°C overnight. DNA was cleaned up by QIAquick PCR purification column (Qiagen) according to the manufacturer’s instructions and eluted in 30 μl of elution buffer (EB).

### ChIP-seq data analysis

The purified ChIPed DNA was sent to IGA (Institute of Applied Genomics, Udine) which performed quality and quantity assessment, library preparation and sequencing using standard Illumina TruSeq for HiSeq 2000 reagents. Each sample was sequenced in single-end (50 bp) mode for a total of 40 million reads per sample. Preliminary sequencing quality assessment was performed using FastQC (http://www.bioinformatics.babraham.ac.uk/projects/fastqc/). The samples passing the literature quality standards were aligned on the human genome (GRCh37/hg19) using BWA using default parameters. In order to maintain the collinearity between the read signal and the protein occupancy on the genome, multiple-matching reads were eliminated using ad-hoc SAMtools (Li et al., 2009) and UNIX shell integrated scripts. Subsequently, peak calling was performed with MACS 1.4.2 (Zhang et al., 2008). Preliminarily, a parameter evaluation step was performed in order to converge on the optimal parameters for the peak calling. Main parameters in MACS algorithm are the MFOLD (enriched quality interval) and the bandwidth (shifting model length). Automatic MACS runs were submitted incrementing both parameters in order to select the most reliable values according to the peak discovery rate. This evaluation was carried out using 2 different proteins characterized by different peak shape (γH2AX and XRCC4). After that, all the proteins underwent peak calling. The output was intersected with the AsiSI sites database using BEDtools (Quinlan and Hall, 2010). In the end, quantitative analysis of induced (+4OHT) and uninduced (-4OHT) dataset was carried out via PscanChIP (Zambelli et al., 2013) using default parameters. In particular, we focused on a proximal region surrounding the AsSI site by tailoring empirically the frame of the algorithm on the peak-length coming from MACS. Moreover, PscanChIP produced a final list of AsiSI sites ranked for the imbalance of γH2AX signal via χ^2^ test. Finally, data visualization was obtained with ngs.plot package, an R based data mining and visualization tool for NGS data (Shen et al., 2014). This tool is based on two steps of normalization, in the first step of length normalization regions of variable sizes are equalized. In the second step, the vectors are normalized against the corresponding library size to generate the so called reads per million mapped reads (RPM) values that allow two NGS samples to be compared regardless of differences in sequencing depth.

### qPCR analysis

Same volumes of immunoprecipitated chromatin were used for standard qPCR on a region proximal to the DSB I or DSB II or a region far from any annotated AsiSI site as control as in (Iacovoni et al., 2010).

Values for each immunoprecipitated sample were normalized on their inputs. qPCR was performed with QuantiTectTM SYBR Green PCR Master Mix (QUIAGEN) on a Roche LightCycler 480. The program used is the following:

1. Denaturation: 95°C 15 min, 1 cycle
2. Denaturation/Annealing/Extension: 95°C 15 sec > 60°C 20 sec > 72°C 30 sec, 50 cycles
3. Melting curve: 40°C > 90°C > 40°C, 1 cycle

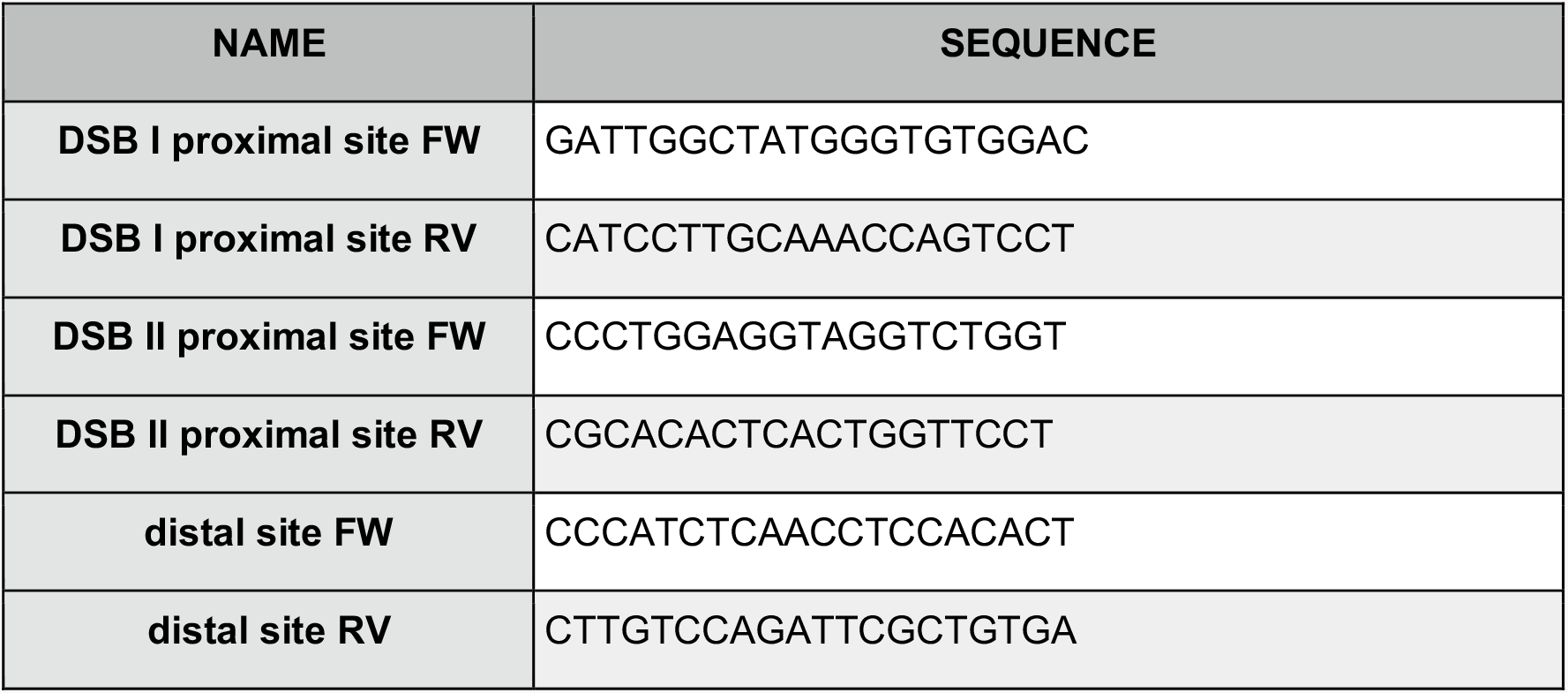

### RNA interference

The ON-TARGET plus siRNA oligonucleotides (Dharmacon) were transfected at the final concentration of 10 nM by Lipofectamine RNAiMax (Life Technologies) following the manufacturer’s protocol. 72 h later, DNA damage was induced and samples were collected. The sequences (5’-3’ orientation) of the siRNA oligonucleotides used are reported in the table below.

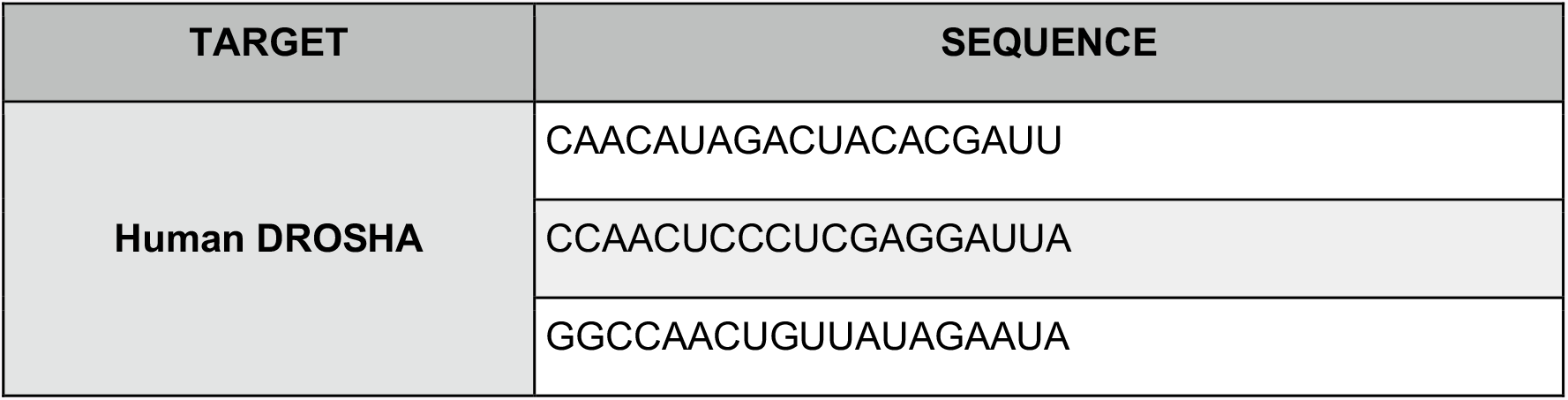
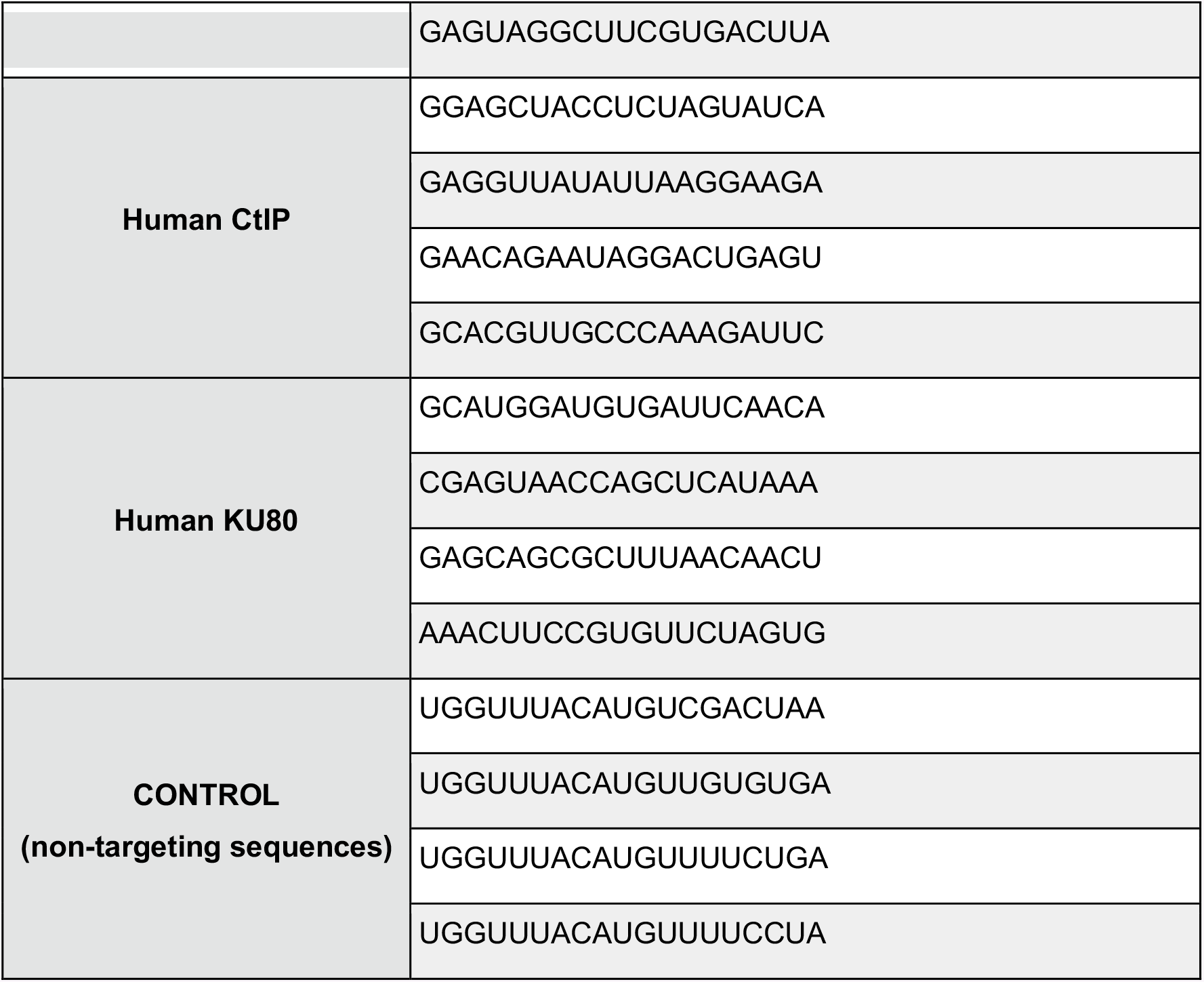

### Indirect immunofluorescence and imaging analysis

Cells were grown on coverslips, fixed in 4% PFA for 10 min at room temperature and permeabilized in Triton 0.2% in 1X PBS for 10 min at room temperature. After 2X washings in 1X PBS, coverslips were blocked in 1X PBG (stock PBG 10X: 5% BSA; 2% gelatin from cold water fish skin in PBS) for 1 h at room temperature. Primary antibody (see Antibodies section for a complete list) incubation was performed for 1 h at room temperature in a humid chamber. After 3X washings in 1X PBG, secondary antibody incubation was performed for 1 h at room temperature in a humid chamber. After 2X washings in 1X PBG and 2X washings in 1X PBS, incubation with DAPI 0.2 μg/mL (Sigma-Aldrich) was performed for 2 min at room temperature. After 2X washings in 1X PBS and 1X washing in deionized water, coverslips were mounted onto glass slides with Aqua Poly/Mount (Polysciences) and let dry overnight at room temperature.

Images were acquired using a widefield epifluorescent microscope (Olympus IX71) equipped with 60X objective. Photomicrographs were taken with digital camera Cool SNAPES (Photometrics). Data acquisition was done using the MetaMorph software (Universal Imaging Corporation). Comparative immunofluorescence analyses were performed using the automated image-analysis software CellProfiler 2.1.1. (Carpenter et al., 2006).

### In situ proximity ligation assay (PLA)

The Duolink^®^ In Situ Orange Starter Kit Mouse/Rabbit (Sigma-Aldrich) was used according to the manufacturer’s protocol. Briefly, cells were fixed, permeabilized and blocked as described above for immunofluorescence studies. Then, cells were incubated with primary antibodies diluted in 1X PBG for 1 h at room temperature (see Antibodies section for a complete list). The cells were washed 3X in 1X PBG and incubated with the PLA probes (secondary antibodies conjugated with oligonucleotides) for 1 h at 37°C in a humid chamber. Cells were washed 2X in Buffer A (supplied with the kit) and the ligation reaction was carried out at 37°C for 30 min in a humid chamber followed by wash in Buffer A. The cells were then incubated with the amplification mix for 1.5 h at 37°C in a darkened humidified chamber. After washing with Buffer B (supplied with the kit) followed by a 1 min wash with 0.01X Buffer B the cells were incubated with DAPI 0.2 μg/mL (Sigma-Aldrich) and mounted.

Images were acquired using a widefield epifluorescent microscope (Olympus IX71) equipped with 60X objective. Quantification of nuclear PLA dots was performed with the automated image-analysis software CellProfiler 2.1.1(Carpenter et al., 2006).

### DNA Damage In situ ligation Proximity Ligation Assay (DIPLA)

After fixation and permeabilization as described for immunofluorescence, cells were treated for DIPLA. Coverslips were washed twice for 5 min in 1X Cut Smart buffer (NEB) and once in 1X blunting buffer (NEB). Afterwards, blunting was performed at room temperature for 60 min, in a final volume of 50 μL for each coverslip using: 38.5 μL H2O, 5 μL 10X Blunting Buffer (NEB), 5 μL dNTP 1mM (NEB), 0.5 μL BSA Molecular Biology Grade 20mg/mL (NEB), 1 μL Blunting Enzyme Mix (NEB). Coverslips were then washed twice in 1X CutSmart buffer and twice in 1X T4 Ligase buffer (NEB). Then in situ ligation was performed overnight at 16°C in a sealed humid chamber, in 100 μL final volume per coverslip using: 2 μL T4 Ligase (NEB), 5 μL 10 uM biotinylated linker (5′TACTACCTCGAGAGTTACGCTAGGGATAACAGGGTAATATAGTTT[biotdT]TTTCTATATT ACCCTGTTATCCCTAGCGTAACTCTCGAGGTAGTA3′), 10 μL 10X T4 Ligase Buffer (NEB), 1 μL dATP solution 100mM (NEB), 1 μL BSA Molecular Biology Grade 20 mg/mL (NEB), 81 μL H2O. Coverslips were washed twice in PBS and processed as described for PLA using a primary antibody against Biotin partnered with a primary antibody directed against the protein under investigation (see Antibodies section for a complete list) (Galbiati et al., 2017).

### Immunoblotting

Cells were lysed in Laemmli sample buffer (2% sodium dodecyl sulphate (SDS), 5% glycerol, 1.5% Dithiothreitol (DTT), 0.01% bromophenol blue, 60 mM Tris HCl pH 6.8). Collected cells were sonicated (Diagenode) with 3 bursts of 15 sec and heated for 10 min at 95°C. 10-15 μl of lysate were loaded on a SDS-polyacrylamide gel with a width of 1 mm along with 7 μl of molecular weight markers (Biorad). A voltage of 60V for the stacking gel and of 150V for the resolving gel were applied. Gels were run in Tris-Glycine electrophoresis buffer (25 mM Tris; 250 mM Glycine; 0.1% SDS). For Western blotting analysis proteins were transferred to a 0.2 μm nitrocellulose membrane (Biorad Trans-Blot^®^ TurboTM transfer pack) using the Trans-Blot^®^ TurboTM Transfer System apparatus (Biorad). The transfer was performed at 25V for 7 or 10 min (according to the molecular weight of the proteins under investigation). Membranes were incubated with 5% skim milk in TBS-T buffer (Tween20 0.1%) for 1 h, followed by over-night incubation at 4°C with primary antibody and 3X washed with TBS-T before 1h incubation at room temperature with the specific HRP-conjugated secondary antibody. Chemiluminescence detection was done by incubation with LuminataTM Classico or Crescendo (Millipore). Proteins were visualized by autoradiography on ECL films (Amersham), using various exposure times and manually developed.

### Cell-cycle analyses by FACS

U2OS cells were collected 72 h post transfection with siRNA, washed in PBS and fixed in 75% ethanol overnight at 4°C. 106 fixed cells for each condition were washed once in PBS 1% BSA and re-suspended in PBS containing propidium iodide (50 μg/ml) and RNaseA (250 μg/ml), and incubated overnight in the dark. FACS analysis was performed on single cell suspensions. For each measurement, at least 10,000 cells were acquired. Samples were acquired either on a FACSCanto II (Becton Dickinson) or a Bio-rad S3e. Propidium iodide was excited with a 488 nm laser and emission was detected with a 670LPnm filter. Data were acquired with FACSDiva 6.1.1 (Becton Dickinson) or Prosort 1.5 (Bio-rad) and analyzed with ModFit LT 3.0 (Verity Software House) or FCS Express 5 (DeNovo) software.

### NHEJ repair reporter assays

The U2OS EJ5 cell line (gift from Dr. Jeremy Stark and prof. Maria Jasin-Memorian Sloan Kettering Cancer Center, NY) was used for assaying NHEJ repair efficiency. Cells were transfected with a plasmid expressing I-SceI or mock transfected with an empty vector for 48h. GFP-positive cells were identified and quantified by flow cytometry. The repair efficiency was scored as the percentage of GFP-positive cells. To examine the role of individual genes in DSB repair, prior to the transfection with I-SceI, cells were treated with siRNAs specifically targeting each gene for 120h.

### Statistical analyses

Results are shown as means plus/minus standard error of the mean (SEM). P-values were calculated by Student’s t-test or one-way ANOVA with multiple comparison. ^*^ indicates p-value<0.05, ^**^ indicates p-value<0.01, ^***^ indicates p-value<0.001, ^****^ indicates p-value<0.0001, according to GraphPad Prism’s statistics. ‘n’ stands for number of independent biological experiments. The number of cells in each experiment is specified in the corresponding figure legend.

## Author contributions

M.R performed ChIP-seq bioinformatic analyses. F.I. contributed to the initial steps of ChIP-seq data. A.G. provided valuable support for DI-PLA experiments. L.C. and S.S. performed FACS analyses of NHEJ assay in EJ5-GFP cells. M.C. performed all the experiments, contributed to experimental design and to manuscript preparation. F.d’A.d.F conceived the study and edited the manuscript. S.F. planned the project, supervised experimental design and execution and wrote the manuscript.

## Acknowledgments

We thank J. Stark and R. Greenberg, for sharing reagents and G. Legube for sharing reagents and unpublished results. We thank M. Cinquanta for the identification of available DROSHA antibodies. We thank F.d’A.d.F. and S.F. group members for support and discussions. M.R. was supported by Fondazione Buzzati-Traverso. S.S is supported by Associazione Italiana per la Ricerca sul Cancro (Start-up Grant 12710) and the European Commission (PCIG10-GA-2011-303806). F.d’A.d.F. was supported by the Associazione Italiana per la Ricerca sul Cancro, AIRC (application 12971), Human Frontier Science Program (contract RGP 0014/2012), Cariplo Foundation (grant 2010.0818 and 2014-0812), Fondazione Telethon (GGP12059), Association for International Cancer Research (AICR710 Worldwide Cancer Research Rif. N. 14-1331), Progetti di Ricerca di Interesse Nazionale (PRIN) 2010–2011 e 2015, the Italian Ministry of Education Universities and Research Progetto Bandiera Epigenomica-EPIGEN, AMANDA projects Accordo Quadro Regione Lombardia–CNR and European Research Council advanced grant (322726) S.F. was supported by Collegio Ghislieri and Fondazione Cariplo (Grant rif. 2014-1215). S.F. and F.d’A.d.F are supported by AriSLA (project “DDRNA and ALS”).

